# Beyond Scenarios - Optimization of breeding program design (MoBPSopti)

**DOI:** 10.1101/2023.04.03.535337

**Authors:** Azadeh Hassanpour, Johannes Geibel, Henner Simianer, Torsten Pook

**Affiliations:** University of Goettingen, Department of Animal Sciences, Center for Integrated Breeding Research, Animal Breeding and Genetics Group, Albrecht-Thaer-Weg 3, 37075, Goettingen, Germany; Institute of Farm Animal Genetics, Friedrich-Loeffler-Institut, 31535 Neustadt, Germany; Wageningen University & Research, Animal Breeding and Genomics, P.O. Box 338, 6700 AH Wageningen, Netherlands

**Keywords:** optimization, resource allocation, kernel regression, genetic gain, inbreeding

## Abstract

In recent years, breeding programs have become increasingly larger and more structurally complex, with various highly interdependent parameters and contrasting breeding goals. Therefore, resource allocation in a breeding program has become more complex, and the derivation of an optimal breeding strategy has become more and more challenging. As a result, it is a common practice to reduce the optimization problem to a set of scenarios that are only changed in a few parameters and, in turn, can be deeply analyzed in detail. This paper aims to provide a framework for the numerical optimization of breeding programs beyond just comparing scenarios. For this, we first determine the space of potential breeding programs that is only limited by basic constraints like the budget and housing capacities. Subsequently, the goal is to identify the optimal breeding program by finding the parametrization that maximizes the target function, as a combination of the different breeding goals. To assess the value of the target function for a parametrization, we propose the use of stochastic simulations and the subsequent use of a kernel regression method to cope with the stochasticity of simulation outcomes. This procedure is performed iteratively to narrow down the most promising areas of the search space and perform more and more simulations in these areas of interest. The developed concept was applied to a dairy cattle program with a target function aiming at genetic gain and genetic diversity conservation limited by budget constraints.

## Introduction

The process of designing a breeding program is a highly complex task. It requires breeders to account for various highly interdependent breeding objectives while considering practical considerations such as budget constraints and housing capacity. While there are tools available, such as the breeder’s equation (Falconer and Mackay 1996), for predicting the outcomes of a breeding program, comparing various breeding schemes with different objectives can be quite difficult, where even a small alteration in one parameter can affect multiple aspects of the program (Henryon et al. 2014; Simianer et al. 2021).

The problem of how to best allocate resources has been addressed in multiple studies, each proposing different optimization strategies ranging from the distinctive features of various mating decisions to the population size, breeding cycle duration, the number of selected individuals, and selection steps (Henryon et al. 2014; Hickey et al. 2014; Woolliams et al. 2015; Gorjanc and Hickey 2018; Moeinizade et al. 2019; Wellmann 2019; Allier et al. 2020; Duenk et al. 2021; Ojeda-Marín et al. 2021; Moeinizade et al. 2022). However, although different aspects of the breeder’s equation include guidance on the use of quantitative genetic theory, and it is a prerequisite for the effective design of new breeding programs, exact formulas to calculate expected gains are usually developed for simple settings with random mating, only one breeding cycle, and a single trait. While these formulas may offer practical guidance for breeding program optimization, their lack of generalizability restricts their usefulness for modern breeding programs that involve many highly interdependent parameters.

Stochastic simulations of breeding programs have been proposed as a useful tool in recent years, with various software options available (Sargolzaei and Schenkel 2009; Faux *et al*. 2016; Liu *et al*. 2018; Pook *et al*. 2020) to execute such simulations for specific breeding actions. Instead of considering cohorts of individuals like in the traditional gene-flow model (Hill 1974), stochastic simulations simulate individuals and all breeding actions, which allow for more flexibility and in-depth modeling of modern breeding schemes.

As the output of a stochastic simulation of breeding programs is the realization of a stochastic process, this provides additional challenges but also opportunities. On the one hand, it is impossible to directly derive the expected outcome/value of key metrics of the breeding scheme (e.g., as done with the breeder’s equation). On the other hand, the main benefit of using stochastic simulations is that they allow for taking into account the inherent variability and uncertainty that exists in any breeding program, and thus access to the associated risks and probabilities to reach certain breeding objectives. Nevertheless, as the expected outcome of a breeding scheme can not be calculated deterministically, optimization has to cope with the additional challenge of stochasticity (Amaran et al. 2016).

Moreover, stochastic simulations require high computing power, as every individual meiosis and every breeding action such as the breeding values estimation is explicitly simulated. Combined with the high number of parameters to consider, it becomes computationally challenging to simulate each potential breeding scheme and directly derive the optimal one. Therefore, to overcome these computational challenges, analysis of breeding programs via stochastic simulations is usually limited to a couple of potentially interesting scenarios, which are then simulated and compared against each other (Wensch-Dorendorf et al. 2011; Esfandyari et al. 2015; Gaynor et al. 2017; Büttgen *et al*. 2020; Pook et al. 2021).

The grid-search approach is a commonly used algorithm for finding the optimum allocation of test resources when there are few parameters to optimize (Longin et al. 2006; Gordillo and Geiger 2008; Mi et al. 2014, 2016; Pook et al. 2021). However, although grid-search provides an acceptable solution to the problem in many smaller applications, as it is relatively straight-forward to define a grid of possible parameter combinations and evaluate the performance of each combination, it is not efficient for a large number of parameters due to the increased computational time and, or the need for a sparse grid.

Our study aims to provide a general optimization framework for breeding programs beyond grid-search algorithms or plain scenario comparisons under a given target function to maximize. Particular focus is on defining the space of all possible breeding programs under constraints like the budget and the contribution of this paper is to provide a framework to cope with the variability in the outcome of stochastic simulations.

## Materials and methods

### General pipeline for optimizing breeding programs

We will propose a general pipeline for optimizing breeding programs in the following. A schematic overview of the different steps is given in Figure 1. Subsequently, we will discuss the individual steps of the pipeline in more detail and discuss them along a classical dairy cattle scheme as a toy example with a detailed description in Supplementary File S1.

**Figure 1.**
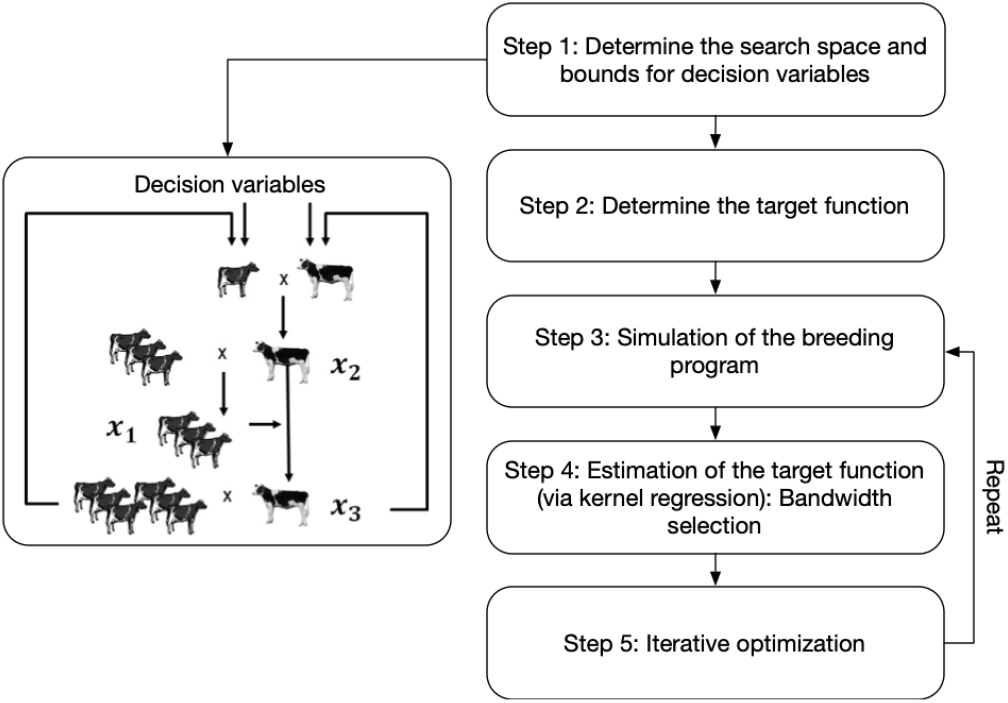
Procedure proposed for optimization via simulation for a breeding strategy.

#### Step 1: Determine the search space and bounds for decision variables

In the case of the dairy cattle example, we want to consider three variables *x* = (*x*_1_, *x*_2_, *x*_3_) for optimization with *x*_1_ being the number of test daughters, *x*_2_ being the number of test bulls and *x*_3_ as the number of selected sires *x*_3_. As in breeding programs, the number of units must be an integer, since it is not possible to produce a fraction of a unit, we modeled this natural constraint by specifying that the decision variables must be non-negative integers.

The cost of a program in the search space refers to the resources required to execute the program. Here, the budget acts as a boundary on the actions that can be taken to achieve the desired outcome, and it is up to each breeder to consider this amount for completing the program. Applying this to our exemplary scenario, we assumed an arbitrary annual budget of 10M € and housing cost of 3,000 € per bull and 4,000 € per cow. Based on our prior expectations, We aim to focus our search efforts on scenarios of significant interest. To do this, we will only consider breeding schemes where the number of test bulls falls within 100 to 700, and the number of selected sires falls within 3 to 30 as we expected other designs to be less efficient and did not consider them to save computing time. In case the optimal solution is found in a corner solution, these constraints may need to be adjusted and softened to ensure that the best solution will not be missed. Similarly, one could imagine practical limitations like a maximum housing capacity in the stable.

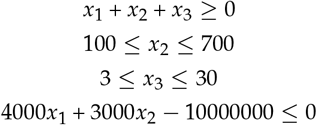

#### Step 2: Determine the target function

To perform any type of optimization, it is required to have a well-defined target function, hence a breeding goal that is a combination of different breeding objectives that are appropriately weighted. In our example, the optimization problem was formulated to achieve two objectives: 1) maximizing genetic gain for the trait of economic importance, and 2) maintaining genetic diversity. With these two objectives, we aim to identify the optimal parametrization, hence the design of the breeding scheme in our exemplary breeding scenario. Our objective function *m* was then created as a composite of the two competing objectives that would allow us to maximize the genetic gain while ensuring that the inbreeding is kept as low as possible. This results in the following:

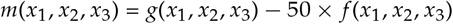

with *g* being the resulting genetic gain and *f* being the inbreeding level after 15 years of breeding. *g* is calculated based on the true underlying genomic values of individuals, and *f* is calculated based on kinship and underlying known points of recombination and founders for each inherited segment. For additional details on kinship calculation, the interested reader can refer to section 9.4 of the MoBPS package user manual (Pook et al. 2022). Note that both are usually not known in practice, but were chosen to represent the underlying true values without estimation errors/biases. In this example, we chose a weighting factor of 50 for *f* to give approximately equal importance to both breeding goals. It’s important to note that this choice was arbitrary and based on the range of values observed in *f* and *g*. The appropriate choice of weighting factor will vary depending on the specific breeding scheme and goals. In the end, in our example, this allows the two objective functions to be compared and allows for a more balanced consideration of their trade-offs.

#### Step 3: Simulation of the breeding program

The process of setting up a simulation script is a crucial component of our algorithm and a determining factor in our pipeline that is often overlooked in scientific manuscripts on simulation studies. In particular, for this application, it is important to write the simulation script in a flexible and general way so that it can handle all parameter settings within the considered search space, which can be particularly tricky when certain cohorts are not generated in each parametrization. In our example, a stochastic simulation of the breeding scheme is proposed using the R package MoBPS (Pook et al. 2020).

In the breeding program described herein, to initialize our pipeline, a large number of simulations need to be performed to obtain a wide coverage of the search space. For this study, we used 60k simulations, which probably is much more than needed and was more or less randomly chosen as the number of simulations finished on our computing cluster in one night. A summary of the simulation process is outlined in Algorithm 1.

**Algorithm 1.**
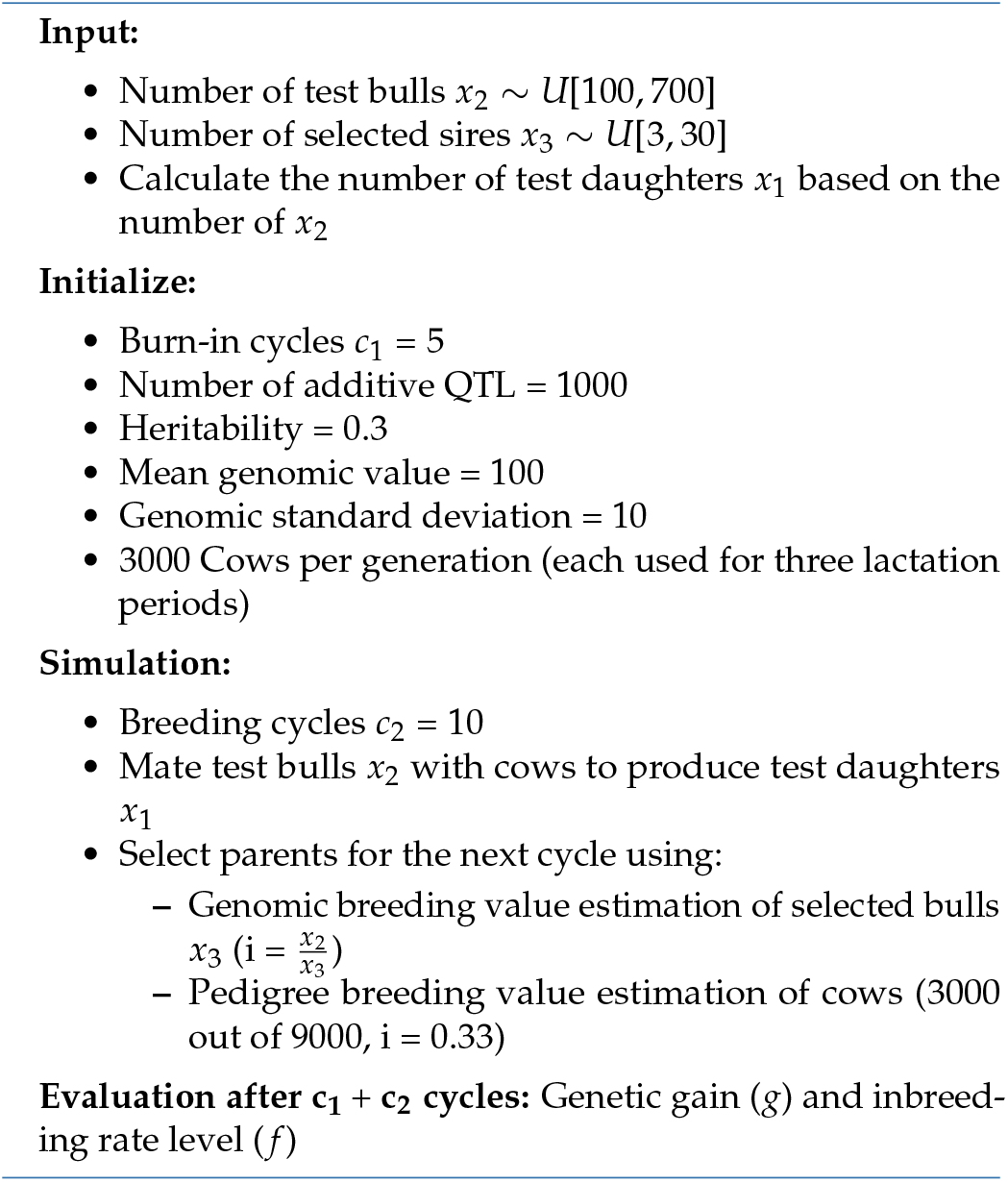
Simulation of Dairy Cattle Breeding Program.

#### Step 4: Estimation of the target function (via kernel regression): Bandwidth selection

The outcomes of our simulation are just realizations of a stochastic process and not calculations of *g, f*, and *m*, respectively. Instead, we want to use these realizations to calculate estimators

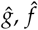, and 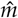. For this, we proposed using kernel regression, via a Nadaraya-Watson estimator (Nadaraya 1964; Watson 1964), which uses a locally weighted average. We can define the estimator of *m* as:

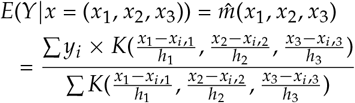

Note that *Y* is a random variable with an expected value *m*(*x*_1_, *x*_2_, *x*_3_) and unknown variance. *y*_*i*_ is the realization of stochastic simulations for our three input parameters (*x*_*i*,1_, *x*_*i*,2_, *x*_*i*,3_). For each value of *x*, the locally weighted average of the *y*_*i*_ is computed with weights given by the used kernel function. We here use a multi-variate Gaussian kernel with independent individual components, resulting in:

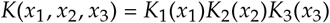

with

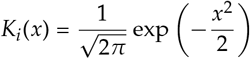

where *x* is the distance between the query point and a data point in the input space, and exp denotes the exponential function. The kernel function is linked to the smoothing parameter bandwidth *h*, which controls the weight given to each simulation in the smoothing procedure. A smaller bandwidth means that the *x*_*i*_ that are closer to *x* will have higher relative weights, and therefore their *y*_*i*_ values will have more influence on our estimate.

It is important to ensure that the bandwidth facilitates the use of appropriate counts of observations at different stages of the estimation process, as there is a well-known bias-variance tradeoff for selecting the bandwidth in high or small-density areas of search space. This can have a significant impact on the accuracy and reliability of the smoothing process, as it determines the shape and width of the smoothing window. A wider *h* will result in a smoother curve with less detail, risk of systematical bias, and oversmoothing. By contrast, a narrower *h* will produce a more detailed curve with more variability. Based on the results in the downstream application, as the inbreeding level changed substantially for a small number of selected sires, kernel regression with large bandwidth can lead to substantial biases. To negate this, the initial bandwidth for *g* (*h*_1_ = 30, *h*_2_ = 30, *h*_3_ = 1) was reduced three-fold for the inbreeding.

### Step 5: Iterative optimization

Results from the kernel regression provide an estimated maximum of the target function. However, kernel regression has both bias and variance that can be reduced by lowering the band-width and more simulations, respectively. For this, we suggest an iterative procedure to use results from Step 4 to identify the most promising areas of the search space for further exploration and then repeat steps 3 and 4, while only performing additional simulations in those areas of interest to reduce the uncertainty in the estimate in those areas.

As the iteration progresses, the smaller bandwidth can capture finer details in the data for fine-tuning. This can help to improve the accuracy of the model by capturing small, localized patterns that may be important for making accurate predictions. The process should then be repeated until the estimated maximum in those areas has reached a stable point and is not likely to change significantly with further exploration, i.e., convergence has been achieved.

## Exploring the effect of variability in stochastic simulation

Due to the inherent randomness of the stochastic simulation process, the estimates obtained from a single stochastic simulation may vary significantly from one simulation to another. To account for this and decrease the variance, it is often necessary to conduct many simulations. This ensures that the results obtained are robust and reliable and that the average of simulations will converge to the true underlying value of the variable we are trying to estimate.

We employed a two-step analysis process to evaluate the effectiveness of addressing variation in our estimate caused by multiple runs of simulation or slight variations in parameter values across simulations. The first step involved calculating the variance of the target function after it had been stimulated. This allowed us to establish a baseline understanding of the level of variability present within the composite function. In the second step, we applied the kernel smoothing method to the target function in order to smooth and reduce the variability of the target function. After applying the kernel smoothing method, we again calculated the target function’s variance. By comparing the variance of the target function before and after the kernel smoothing method, we can determine the smoothing process’s effectiveness in reducing the target function’s variability. We compared the variance of the target function within distinct three-dimensional windows before and after applying the kernel estimator.

Furthermore, we aim to evaluate the precision of determining the target value within one iteration by determining the appropriate number of simulations required. Further details of this method can be found in Supplementary File S2.

## Results

In the first step with 60k simulations, the genetic gain is maximized when using the lowest number of selected sires, combined with the highest number of test daughters (2425,100,3), which results in a genetic gain of 12.7 with a genetic standard deviation of 1.04 after ten breeding cycles (Figure 2a). Here only the two dimensions based on *x*_2_ and *x*_3_ were referred to, as the number of test daughters (*x*_1_) depends directly on the number of test bulls (*x*_2_).

**Figure 2.**
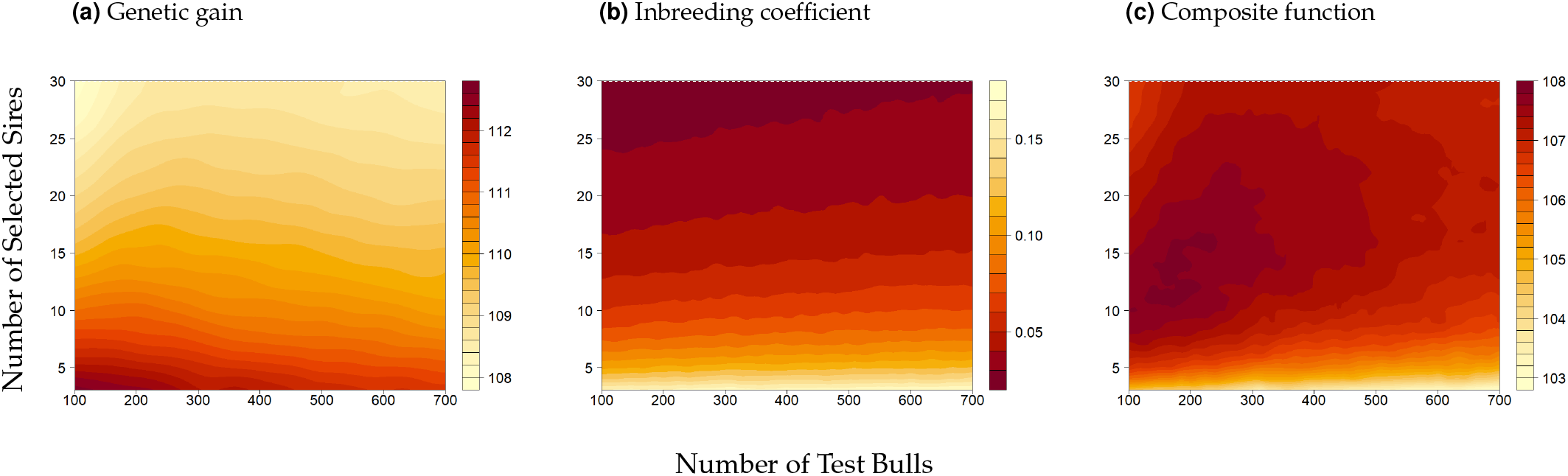
Visualization for Nadaraya-watson estimator for (2a): Genetic gain (*ĝ*) with a bandwidth (*h*_2_ = 30, *h*_3_ = 1), (2b): Inbreeding coefficient 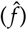 with a bandwidth of *ĝ/*3:, (2c): Composite function 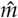 based on 60k simulations. The colors represent the relative value of the target function, with dark red showing the favorable outcome of the target function.

In the example of our dairy cattle breeding, lower selection intensity and increasing the number of test daughters can lead to more accurate predictions of breeding values. This is because more test daughters provide more data to estimate the breeding values of the sires. On the other hand, increasing selection intensity can be obtained either by decreasing the number of selected bulls or increasing the number of test bulls, which, however, conflicts with the number of daughters tested. As Figure 2a shows the benefit of having more accurate predictions of breeding values through more test daughters outweighs the potential increase in selection intensity in our toy scenario. Note that if only optimization of *g* would be the target, the initial constraints should be less strict, allowing the optimization algorithm more freedom to find a better solution by investigating a lower number of test bulls and selected sires.

On the contrary, lowering the intensity of selected sires leads to an increase in genetic diversity, allowing for a higher range of genetic variation to be maintained. The result is shown in Figure 2b where the inbreeding was minimized using the highest number of selected sires, combined with the highest number of test daughters (2422, 103, 30), which resulted in an inbreeding coefficient of 0.025.

In a contour map shown in Figure 2b, the distance between contour lines represents the steepness of the rate of change of the inbreeding function, where for a small number of selected sires, the inbreeding level changed substantially. This suggests that using a smaller bandwidth for the inbreeding function, in comparison to the genetic gain, resulted in a more accurate, smoother fit. This is because the overall variance observed in the inbreeding outcome was relatively low, and a smaller bandwidth was able to capture this variation more effectively.

Combining both objective functions, the estimated maximum of the composite function 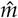 based on the kernel estimator is obtained at (2332,224,16) with the best value of 107.8528 (Figure 2c). Note, however, that the zoom-in plot of the composite function shows three local optima, where two different local maxima with similar values (2353,189,12) and (2371,172,14) lead to a genetic gain of 107.8436 and 107.8415, respectively. Thus, this first iteration is not sufficient to narrow down the absolute maximum. However, it allows us to narrow down the search space for subsequent iterations to 120 ≤ *x*_2_ ≤ 260 and 10 ≤ *x*_3_ ≤ 20, as all three values fall within this area (3a).

As a result, an additional 50k simulations were conducted in the second iteration, this time focusing only on the promising areas of the search space (areas inside of the black square in Figure 3b). In the second iteration, the search space size was significantly reduced by narrowing the grid from a 27×600 grid to a 10×140 grid, resulting in an 11-fold decrease in the search space.

**Figure 3.**
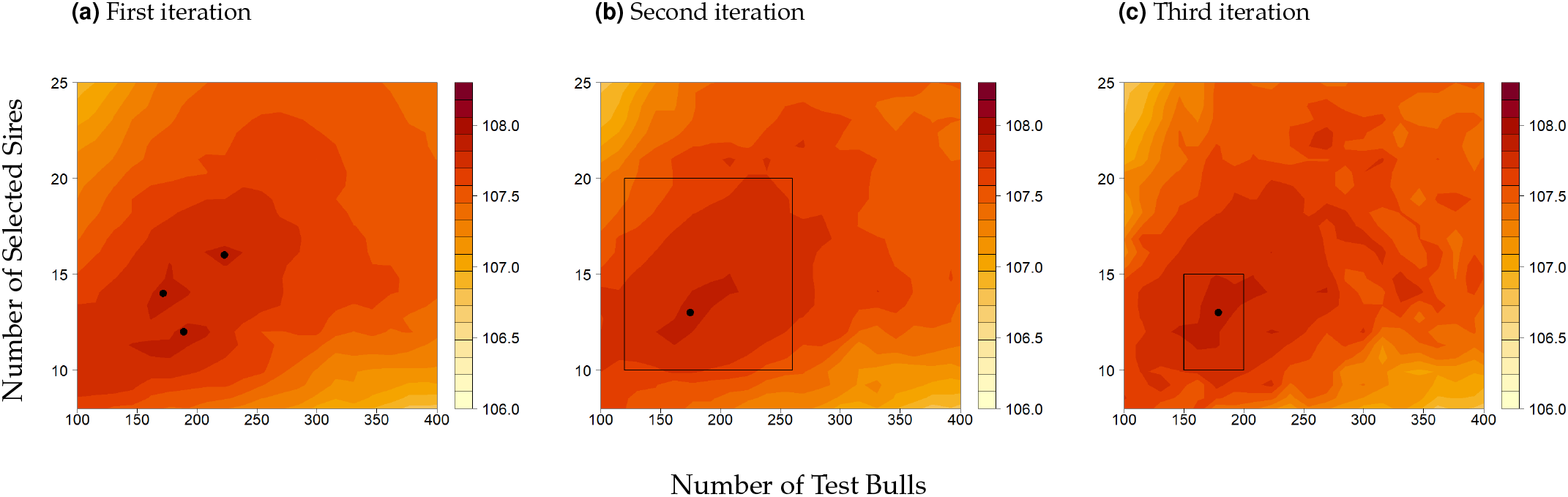
Smoothed surface contour map for the composite function of (3a) full space (zoom-in plot of 2c to show three local optima): First iteration with a bandwidth (*h*_2_ = 30, *h*_3_ = 1), (3b) Second iteration with a bandwidth (*h*_2_ = 15, *h*_3_ = 0.5) (3c) Third iteration with a bandwidth (*h*_2_ = 7.5, *h*_3_ = 0.25). The indicated bandwidth here refers to the bandwidth for *g*, and this bandwidth for *f* in all iterations was divided by three. Dots indicate local maxima and the area inside of the squares indicates the search space in the current iteration. The area outside the square shows the prominent effect of the bandwidth and the variance of our estimates, which is related to the number of simulations used to estimate the kernel function.

As the number of simulations increased within the defined window size for calculating the kernel function, the variance decreased compared to the first iteration. This allows us to reduce the bandwidth, allowing for a better approximation of our composite function and a smoother curve. Therefore, the bandwidth, which controls the degree of smoothing for the second iteration, was halved. This results in the best solution 107.89 from optimization, suggesting (2368, 175, 13) as the optimum.

In the third iteration, we conducted an additional 12k simulations by reducing the grid from 10×140 to 50×5, where the size of the search space was decreased by 5-fold. The reduction in the search space size enabled an increase in the number of observations within the defined window and a reduction in bandwidth by half. The optimization suggests (2365, 179, 13) indicating stabilization of the optima with the best value of 107.91 (Figure 3c).

Note that as no further simulations were run outside of the areas of interest, the chosen bandwidth will be too small to reliability estimate *m* outside of the new search space, e.g., as seen in Figure 3c, there are various local extrema, however, as they are all substantially lower than the estimated global optimum, they can be safely ignored. If a specific region shows high potential, it may be worthwhile to run additional simulations in those previously disregarded regions. This will be particularly relevant for more complex optimization problems with more parameters.

By comparing the variance of the target function before and after the kernel smoothing method in the first iteration, we determined how much the smoothing process has reduced the variability of the target function. A substantial decrease in variance in Figure S 1 indicates that the kernel smoothing method is performing well and effectively reducing the overall variability of the target function, leading to more accurate results due to having less uncertainty associated with our estimates and predictions.

Identifying the global maxima and, or the target area for further iterations based on the simulation of the first iteration has a significant amount of variance. When just conducting 1000 simulations, the estimated global maximum for *m* is estimated to be in a corner solution in 11% of all runs with 100 ≤ *x*_2_ ≤ 120 (Figure S 2a) and some estimated optima with *x*_2_ value of up to 380. Only 16% of all estimated global maxima fall within the range (150 ≤ *x*_2_ ≤ 200 and 10 ≤ *x*_3_ ≤ 15) that we chose for final investigation. In comparison 10k (S 2b), 20k (S 2c) and 60k (S 2d) simulation result in 30%, 38%, 47% of the runs in our afterward chosen search space.

### Computing time for simulation

A server cluster with Intel Platinum 9242 (2×48 core 2.3 GHz) processors was used for this study. Simulations were performed on the single core and required ∼15 minutes and 15GB RAM peak memory usage per simulation.

## Discussion

In this study, we have developed a general optimization framework for breeding programs that goes beyond the limitations of traditional methods, including accounting for the variability in the outcome of stochastic simulations. Insights from the results highlight five key points for discussion:

### Kernel estimator vs grid search/scenario testing

The algorithm discussed in this study offers additional benefits over traditional methods like grid search and plain scenario comparison. Traditional methods usually only consider a limited number of scenarios. In contrast, our algorithm can consider a much more extensive space of scenarios, increasing the chances of finding the optimal scenario for a resource-limited breeding program using stochastic simulations.

The larger search space allows for a more thorough exploration of the problem, which can provide a deeper understanding of the underlying relationships and dependencies between different parametrization. This can be valuable for identifying trends or patterns that may not be immediately visible with a smaller search space. Our algorithm also uses a more efficient search strategy, quickly narrowing down the range of possibilities, and allowing for a more targeted exploration of the parameter space without testing all scenarios. This makes it a more efficient and effective alternative to grid search, which requires many scenarios to be considered to cover all possible combinations of parameters. E.g., imaging that the optimum for a parameter is 150, and if we use a grid-search algorithm that only considers a limited set of discrete values, such as 100, 200, 300, and 400, we miss the true optimum and settle for a suboptimal value of 100 or 200 as the best solution, which could lead to significant performance degradation compared to the optimal setting of 150.

It is important to note that the size of needed simulations required to optimize a model increases exponentially with each new parameter added, making the use of kernel regression challenging to produce such a large amount of simulations to obtain reliable estimates for each region of search space (Lavergne and Patilea 2008; Geenens 2011). Therefore, it is crucial to weigh the benefits of adding additional parameters against the potential limitations they can impose on the accuracy and generalization of the model. Adding too many parameters can lead to overfitting and reduced model performance, as it increases both the computational complexity of the algorithm and the resources required to run it.

### The impact of bandwidth

A local smoothing method is a powerful tool for making predictions based on simulated data, but it can be sensitive to the choice of bandwidth. If the bandwidth is too large, it will assign too much weight to parametrizations far from the target value, which can introduce prediction bias. On the other hand, if the selected bandwidth is too small, there will be a low number of observations, which can lead to high prediction variance. Thus, an appropriately chosen bandwidth will weigh between those two factors. In our particular case with three variables, fitting the bandwidth via visual inspection was sufficient, which facilitates our understanding of the bandwidth behavior.

However, with many parameters, visual and manual bandwidth selection can become difficult. For this, various automated methods for bandwidth selection, such as cross-validation (Hardle and Marron 1985; Jones et al. 1996) and variance-based approaches (Brockmann et al. 1993), can minimize mean-squared errors and aid in selection. When using such an automated procedure for our example (particularly for the first iteration), the suggested bandwidth was smaller than what we used in this study. In general, we would recommend using conservative choices with a relatively large bandwidth in the early steps as the focus in the early steps is not unbiasedness but the identification of target areas for further testing.

### Target function

Formulating an adequate target function is an important part of any decision-making breeding process, which requires careful consideration of both short and long-term objectives, as well as weighing the risks associated with each option, which plays an important role in determining the optimal solution of the optimization problem. Inappropriate weighting between different objectives in the target function can lead to optima that may not be meaningful.

In the example provided, the weighting between *g* and *f* was chosen arbitrarily, but in practice, this process requires more thoughtful consideration due to the limited options available. For example, a company focused on economic production may prioritize traits such as yield over inbreeding and thus may choose a lower weighting factor for inbreeding compared to a conservation breeding program, where the goal is to preserve genetic diversity, and thus the weighting factor for *f* can be higher. Besides, one could imagine a target function that reflects the economic impact of the breeding program, with further potential issues to quantify how much impact an improvement of e.g., fertility trait (in genomic SD units) would lead to how much increase in average annual net profit. This enables breeders to make sure that whatever resources are being spent result in expected outcomes without sacrificing too much money upfront.

Apart from calculating overall economic gains, there is considerable interest in how changes in inbreeding affect the distribution of total gains over many years. In practice, it is common to set a threshold for the maximum amount of inbreeding gain per year to ensure that the population’s genetic diversity is not compromised. In this situation, one could also consider using a target function like our exemplary case that considers genetic gain and inbreeding. This can be done by testing different penalty values placed on two objectives to find the optimal target function that results in inbreeding rates similar to those chosen for the breeding scheme or consider a non-linear weighting of parameters, e.g. applying a high penalty on the target function when a threshold value for a yearly inbreeding rate is exceeded. Finally, these solutions can be analyzed in detail, allowing the breeders to adjust the trade-offs between the different objectives.

### Number of needed simulations

From the distribution of the number of needed simulations results in Figure S2a-S2d, we conclude that, as a general pattern, the more simulations that are run, the more likely it is that the optimal or near-optimal solution will be found. The choice of the number of needed simulations is an important decision to make when designing an iterative simulation-based optimization study. To achieve the best results, adjusting and refining the number of simulations in each iteration is often necessary.

This can be especially tricky when dealing with a large search space that needs to be reduced quickly and efficiently. Using an unnecessarily large number of simulations wastes resources. In contrast, a small number of simulations may produce unreliable results. For basic optimization procedures, choosing fewer simulations might be feasible to obtain the desired statistical power but making such decisions for complex optimization problems with various inputs is not straightforward and it will be highly dependent on the breeding program at hand.

Our findings suggest that achieving good results using a small number of simulations in the first iteration is unlikely (S 2a). In this situation, when an estimator leads to highly different values, it has high variance and is considered less reliable. High variance can result in large fluctuations in the estimator’s value, making it difficult to predict the value of the parameter being estimated accurately. The resulting solutions can be many local optimums, some located in one corner of the parameter space rather than representing an overall optimal result. Overall, this leads to a trade-off between the initial search space is large enough to include all promising settings while not considering settings that are known to be impractical or resulting in lower values in the target function. As the dimensionality of the problem is not as high in our particular example and the shape of 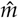 should be relatively smooth, in our case 10k simulations would probably be sufficient. In particular, even if the global optima would not be included in the new search space, the new optimization would most likely end in a corner solution that would signal the need to expand the search space in that particular direction.

Additionally, careful consideration should also be given to external factors such as time, hardware capabilities, or software availability, which cannot be directly altered by users but must still be considered when designing simulation studies. One approach that can be used successfully in this situation is to consider the amount of time one is willing to allocate to the optimization process. How much computing time to spend on which step of the pipeline will be highly dependent on the application but as a rough guideline we would be recommended to use at most 1/3 of the total available computing time in the first iteration. This initial iteration can be beneficial to quickly identify areas of the search space that are likely of low interest and can be disregarded for further investigations.

After that, the number of simulations in subsequent iterations can be increased, however, this should only be done within smaller search spaces that have been identified as promising. This approach then allows for quick progress while also ensuring accuracy by focusing on areas of interest, avoiding running too many simulations in areas where the optima are not likely to be found, and avoiding wasting resources on running simulations that do not yield meaningful results.

Another strategy for optimization is to run simulations and then apply kernel regression to analyze the results. If the results are not smooth enough, running additional simulations used in the kernel regression analysis may be necessary. By continuously analyzing and refining the results, one can improve the optimization process to achieve the best possible outcome in each iteration. It is important to note that while it is true that running a high number of simulations can increase the likelihood of finding an optimal solution, it is important to consider the trade-off between optimization and computing time and there is still randomness in the stochastic simulation outputs. However, sampling error and variance can be reduced by expending additional simulation effort to achieve a predetermined level of statistical power for the optimization strategy. As a result, more simulations provide a high prediction quality in areas close to the optima.

Moreover, the extent of the number of needed simulations varies from objective function to another, and the rate of increase depends on the complexity of the objective function and the number of parameters involved. In general, more simulations may be needed to find the optimal solution if the problem is complex or if the input variables interact in a non-linear way. For our target function with few parameters, increasing the number of needed simulations from 10k to 60k did not result in significant changes to the identification of important optima areas.

### Simulating breeding schemes: The fine line between realism and efficiency

Simulation plays a vital role in understanding the complexities of breeding schemes, but creating an accurate and efficient simulation can be challenging. A realistic simulation must take into account various factors such as genetics, environmental and management conditions, and other relevant considerations. However, including too much detail can make the simulation slow and difficult to run. It is crucial to consider the purpose of the simulation and the level of detail necessary to achieve that purpose and to strike a balance between realism and computational efficiency. Not all factors may be equally important and some details that similarly affect all parametrizations might be worth to be excluded to ensure fast simulation. By making strategic choices about what details to include and what to exclude, the simulation can provide valuable insights while still being efficient to run, where the goal of the simulation should still provide an accurate representation of reality for the intended breeding purpose.

## Conclusion

In conclusion, the Nadaraya-Watson estimator has proven to be a valuable tool in optimizing breeding programs with few parameters. Its flexibility in considering a large space allows for accurate predictions, and its ability to reduce variability in optimization strategies using stochastic simulations provides a more reliable assessment of potential solutions. This promising strategy opens up avenues for further research in optimizing test resources and tackling larger-scale breeding optimization tasks to maintain high accuracy while also considering practical limitations such as limited hardware capacities.

## Data availability

The exact model for the simulation script (File S3), visualization script (File S4), and generated data for reproducing the results have been shared with the scientific community at Figshare: https://figshare.com/s/b30dc547013c9a5a3bd2.

## Acknowledgments

The authors acknowledge the computational support from the Scientific Compute Cluster at GWDG, the joint data center of the Max Planck Society for the Advancement of Science (MPG), and the University of Goettingen.

## Funding

This research was supported by BASF Belgium Coordination Center.

## Conflicts of interest

The authors declare that they have no known competing financial interests or personal relationships that could have appeared to influence the work reported in this paper.

## Supplementary Materials

### File S1

#### Dairy cattle simulation: A classical breeding approach

Briefly, the simulation is designed to imitate a traditional dairy cattle breeding scheme. For this, we are considering selection for ten generations using a single trait with a heritability of 0.30. In this program, as the performance traits of a bull cannot be determined phenotypically and can be expressed only by cows, pre-selected bulls, hence the number of test bulls, *x*_2_ must be mated to cows to produce test daughters *x*_1_. The offspring performance of the test bulls was used as a criterion for selection decisions. The selection of bulls is assumed to be based on genomic breeding value estimation, while cows were selected via pedigree breeding value estimation. The last three cow generations were considered for selection, with phenotypic information available for the two older generations. Bulls were subsequently selected as selected sires *x*_3_ based on their offspring performance. Note that this is a traditional breeding scheme, and particularly on the male side of the breeding scheme generation intervals can be substantially reduced by the use of genomic selection (Schaeffer 2006). To ensure a realistic starting point, the additional five generations were simulated as five burn-in cycles.

### File S2

#### Estimation of the number of needed simulations for optimization

To understand the relationship between the number of simulations we need to run and the accuracy of our optimization process, we used a dataset generation procedure that included smoothing the data and creating different sample sizes from an original dataset (60k simulations). These sample sizes are then used to optimize our objective function by comparing the solution we found using the current sample size (the proposed optimum) to the solution that is expected to be found using a large sample size (the final optimum, which was found in the third iteration).

For this, the kernel density estimator (KDE) was used to calculate a density function for the estimated optimum based on the given number of simulations. It is based on the idea of using a kernel function to smooth out the data and estimate the underlying distribution of the data. This calculation gives us an estimate of how likely it is to miss the maximum by picking a new search space that does not include the optima and how sample size affects the accuracy of our optimization process. The KDE is defined as:

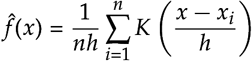

where *f* (*x*) is the estimated density at a given value of *x, n* is the number of data points, *h* is the bandwidth, which controls the smoothing of the data, *x* is the value at which the PDF is being estimated, *x*_*i*_ are the data points, and *K* is the Gaussian kernel function. The consistency of the process was calculated to estimate a stated confidence level with a bootstrap sample of 100.

**Figure S 1.**
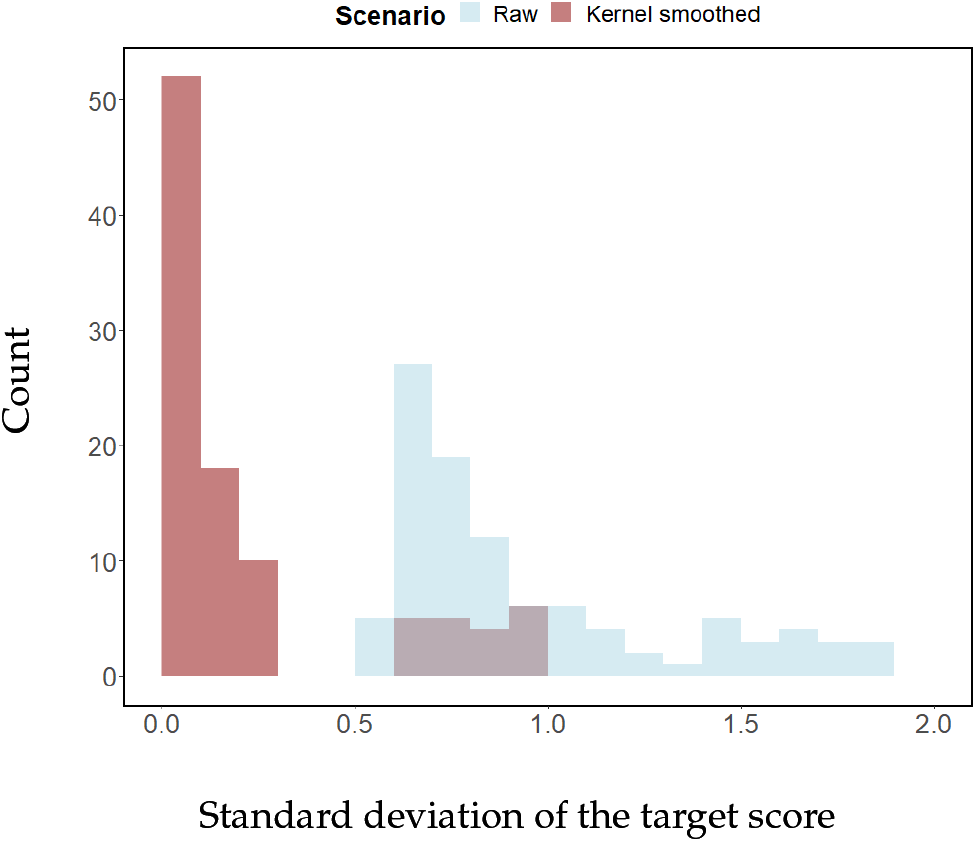
Distribution of the standard deviation of the target function in our search space before and after applying the kernel estimator.

**Figure S 2.**
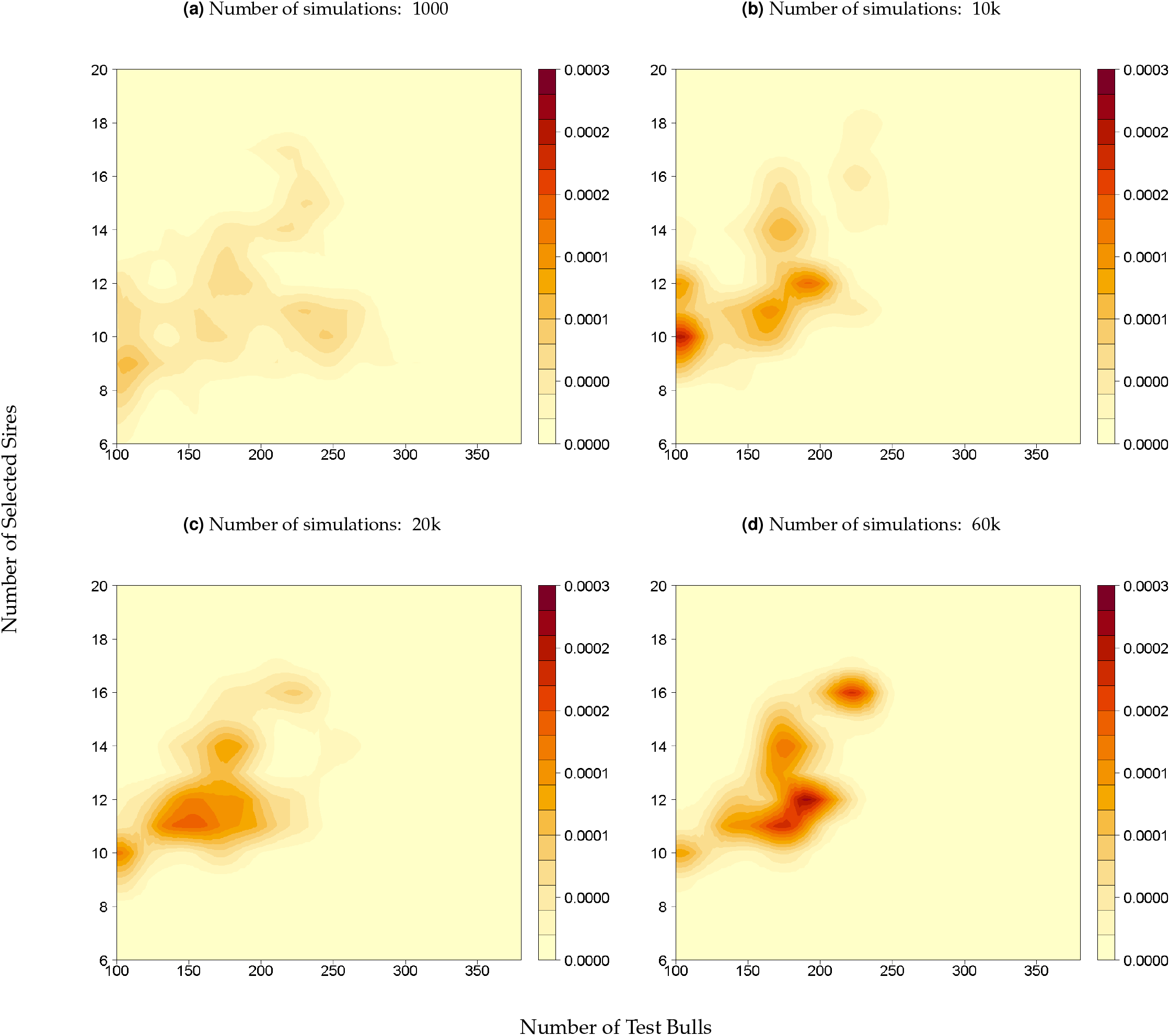
Estimates of the optimum values after smoothing using KDE associated with the number of needed simulations: (S 2a) using 1000 simulations, (S 2b) using 10k simulations, (S 2c) using 20k simulations, (S 2d) using 60k simulations. The spacing between the contours indicates the variation in the data, with areas of higher concentration represented by denser contours and areas of lower concentration represented by more widely spaced contours. The figure shows different shades of color, with darker shades indicating the higher density of the outcome of stochastic simulations with a higher likelihood of finding the global optimum and lighter shades indicating lower density.

## Notes

https://figshare.com/s/b30dc547013c9a5a3bd2

### Competing Interest Statement

The authors have declared no competing interest.

